# CLN5 deficiency impairs glucose uptake in Batten disease

**DOI:** 10.1101/2025.01.02.630719

**Authors:** Maria Marchese, Sara Bernardi, Rachele Vivarelli, Stefano Doccini, Lorenzo Santucci, Asahi Ogi, Rosario Licitra, Jingjing Zang, Rabah Soliymani, Serena Mero, Stephan CF Neuhauss, Giovanni Signore, Maciej M. Lalowski, Filippo M. Santorelli

## Abstract

CLN5 disease, a form of juvenile dementia within the neuronal ceroid lipofuscinosis (NCL), is associated with mutations in the *CLN5* gene encoding the lysosomal bis(monoacylglycero)phosphate (BMP) synthase, essential for BMP production and lysosomal function. Limited knowledge of cellular mechanisms and unclear drug targets hinder translating this to children’s treatment, which remains symptomatic. We developed and characterized a new *cln5* knock-out zebrafish model that replicates key features and molecular signatures of the human disease. Loss of Cln5 function *in vivo* altered axonal growth of retinal ON-bipolar cells revealing new disease features. Additionally, multi-omic analyzes at different developmental stages, revealed an impaired glucose metabolism as an original finding in NCL. This work demonstrates the profound metabolic impact of CLN5 dysfunction, offering a promising avenue toward targeted therapies for juvenile dementia.

## Introduction

The neuronal ceroid-lipofuscinoses (NCLs), also known as Batten disease, are neurometabolic lysosomal storage disorders and collectively represent the most frequent childhood-onset neurodegenerative diseases and the main cause of childhood dementia in Europe [1]. NCLs are characterized clinically by the combination of myoclonus, seizures, blindness, and progressive neurodegeneration leading to dementia. Lacking in most cases specific therapies [2,3], both life expectancy and quality of life are limited in NCL children. The CLN5 disease (MIM: 256731) represents an ultra-rare variant late-infantile form of NCL with onset between 4 and 7 years, caused by biallelic mutations in *CLN5*. CLN5 is highly expressed in the cerebellar Purkinje cells (PC), cortical neurons and hippocampal pyramidal cells, and hippocampal interneurons [4], all areas known to be degenerated in CLN5 human brains [5]. Importantly, hypomorphic gene variants in *CLN5* also occur in other relatively more common genetic conditions such as autosomal recessive macular dystrophy [6], or cerebellar ataxia [7], or even in common neurodegenerative disorders with impaired vision, cognition and movement disorders including Alzheimer’s Disease (AD) and Parkinson’s Disease (PD) [8,9,10]. This implies that research data gathered in the ultra-rare CLN5 can serve to decipher multiple disease conditions. A recent outbreak in the field demonstrated that *CLN5* encodes the lysosomal bis(monoacylglycero)phosphate synthase (BMPS), the enzyme responsible for the production of bis(monoacylglycero)phosphate (BMP; also called lipid lysobisphosphatidic acid, LBPA) [11]. BMPS-deficient cells exhibited a massive accumulation of the BMP synthesis precursor lysophosphatidylglycerol (LPG), depletion of BMP species, and dysfunctional lipid metabolism [11]. BMP is thought to play a role in glycosphingolipid degradation and cholesterol transport, although the precise mechanism of its biosynthesis and transport remains still unclear. In line with the alterations in lipids, the mitochondrial-lysosome interplay, lipid homeostasis and sphingolipid metabolism we had shown in knock-out SH-SY5Y neuronal-like cells [12], new findings on CLN5 function [11] imply that neuronal cells may be especially susceptible to lysosomal disruptions, a feature already known in early biochemical work from cerebral cortex in NCL patients [13]. Interestingly, alterations of BMP levels have been observed in several human diseases [14,15], suggesting that pharmacologically activating and inhibiting the BMP synthase can exert therapeutic effects not only in lysosomal storage disorders such as CLN5 but also in more common forms of dementia [16].

In this light, comprehending metabolic alterations caused by CLN5 impairment during development may constitute a burgeoning research area underlying potential treatments for neurodegenerative disorders, such as juvenile-onset neuronal ceroid lipofuscinoses.

Zebrafish represents a valid model to study the pathomechanisms in NCLs [17,12,18,19,20], and for drug discovery [20]. The main advantage of zebrafish models resides in tackling complex biological problems and offers insight into the cellular processes that influence disease in relatively simpler organisms with high synteny and sequence identity with human genome.

Herewith, we generated a novel *cln5* knockout zebrafish model. Comprehensive behavioral, physiological and molecular characterization of the *cln5*^*−/−*^ model highlights its potential for investigating the *in vivo* mechanisms underlying BMP alteration, and for developing new small-molecule therapies for NCLs.

## Materials and methods

### Zebrafish maintenance

Experiments were conducted on a *cln5*^*−/−*^ and also on a transgenic [*Tg(neurod1:GCaMP6F)*] line (kindly donated by Claire Wyart of the Institut du Cerveau et de la Moelle Épinière, Paris, France) [21] in the nacre (*mitfa*^−/+^) background, which will henceforth be referred to as wild-type (WT). We had previously verified the absence of any behavioral differences between the [*mitfa*^−*/+*^; *Tg (neurod1:GCaMP6F)*] line and the standard AB line [22]. Moreover, for the analysis of retinal bipolar ON-cells we used the transgenic line *Tg(nyx:GAL4-VP16)* (kindly donated by Jeff S. Mumm of the Wilmer Eye Institute, Johns Hopkins University, Baltimore, MD) [23], while for the analysis of Purkinje cell layer we use the line *Tg(aldoca:GAL4FF)* line [24]. The *cln5*^*−/−*^ experimental fish were generated by intercrossing *cln5*^*−/−*^ males and females. Adults were housed in tanks at a density of no more than five zebrafish per liter at a constant temperature of 28.5 °C with a 14-h light/10-h dark cycle. Zebrafish eggs and embryos were collected and grown at 28.5 °C in egg water obtained using “Instant Ocean” sea salts (60 μg/mL) (Aquarium Systems, Sarrebourg, France) and in E3 medium (5mM NaCl, 0.17 mM KCl, 0.33 mM CaCl_2_ 2H_2_O, and 0.33 mM MgSO4, per 1 L of deionized water), respectively, according to established procedures [25, 26]. They were staged in either hour post fertilization (hpf) or days post fertilization (dpf). All compounds used for E3 medium solution preparation were purchased from Sigma-Aldrich (St. Louis, MO). The generation of the mutant, using CRISPR/Cas9 technology, was performed with the ethical approval of the Italian Ministry of Health (approval n° 584/2019-PR), in accordance with the European Union’s Directive 2010/63/EU on the protection of animals used for scientific purposes, under the supervision of the University of Pisa Animal Care and Use Committee, and in compliance with the 3R principles.

### Establishing mutant lines

The selected sgRNA was chosen among the top targets identified by the CHOPCHOP software (www.chochop.rc.fas.harvard.edu/index.php) set with NGG PAM sites and zero predicted off-targets (fewer than three mismatches in the *cln5*-targeting 20-mer). The sgRNA was designed against exon three of the *cln5* transcript (ENSDART00000110866.4) (**Supplementary Table S1**), and generated as already described [20]. The optimized Cas9 mRNA, for genome editing in zebrafish, was transcribed from a linearized template plasmid pCS2-nCas9n using the mMESSAGE mMACHINE^™^ SP6 Transcription Kit (Thermo Fisher Scientific, Waltham, MA). The injection strategy has been performed as already described [20].

### Genotyping

For mutation screening, sgRNA-injected F0 embryos, raised to adulthood and outcrossed with the *Tg(neurod1:GCaMP6F)* line, were used to obtain heterozygous F1 embryos. Adults potentially carrying mutations were identified by PCR and fragment analysis using genomic DNA from 10 randomly selected F1 embryos. Heterozygous F1 fish carrying a 2-bp deletion and missense mutation in the target site were selected and inter-crossed to generate the homozygous F2 *cln5*^*−/−*^ line.

### *In situ* hybridization

Whole-mount *in situ* hybridization was performed in 5 dpf WT larvae as described elsewhere [27].

### Analysis of larval morphology

Live zebrafish at 5 dpf were mounted on glass depression slides with 1% low melting point agarose. Images were obtained using a Leica M205FA stereomicroscope (Leica Microsystems, Wetzlar, Germany). Zebrafish standard body length, eye size, and yolk sac area were measured using Danioscope software (Noldus Information Technology Wageningen, The Netherlands) [22].

### Quantitative (q)RT-PCR

Gene expression levels of *cln5* mRNA used RNA pooled from WT and *cln5*^*−/−*^ larvae. Total RNA was extracted from 30 larvae at 120 hpf using the Quick RNA Miniprep Kit (Zymo Research, Irvine, CA) according to the manufacturer’s instructions. Extraction of cDNA and qRT-PCR were performed as described elsewhere [27]. Relative expression levels of each gene were calculated using the 2^−ΔΔCT^ method [28]. The results obtained in at least three independent experiments were normalized to the expression of the housekeeping gene, *β-actin* (ENSDARG00000037746). The expression analysis of *cln5* mRNA in mutant larvae was calculated setting the mean of the controls at one, and the p-value was calculated using GraphPad Prism 6 software (GraphPad Software Inc., San Diego, CA).

### BMP lipid dosage

In order to evaluate the activity of Cln5 we measured in zebrafish the production of the lipid (18:1/18:1) BMP (S,S) by LC-MS analysis. Lipids were extracted from the *cln5*^*−/−*^ mutant and WT whole brain using a modification of a previously reported method [29]. Briefly, extracted brains were frozen at -80 °C until processed. Next, the sample was put in a 1.5 ml microcentrifuge tube and diluted with 100 μl of 150 mM NaCl aqueous solution and 600 μl of 0.0625 μM DMS (N, N-dimethylsphingosine) in MeOH/CHCl_3_ 1/2. The biphasic solution was incubated at 25 °C for 30 min at 1000 rpm and then centrifuged at 13.000 rpm for 10 min at 4 ^º^C. The lower chloroform phase was recovered, concentrated *in vacuo*, and reconstituted in a MeOH/H2O/i-PrOH 50/45/5 with 0.1% HCOOH and 1 mM ammonium formate and processed immediately. LC-MS analyses were performed in an Orbitrap Exploris 120 mass spectrometer, interfaced with a Vanquish HPLC chromatograph. Analyses were carried out on a Phenomenex Proteo column (150 × 0.3 mm) at a flow rate of 5 μL/min using MeOH/H2O/i-PrOH 50/45/5 with 0.1% HCOOH and 1 mM ammonium formate (phase A) and MeOH/i-PrOH 50/50 with 0.1% HCOOH and 1 mM ammonium formate (phase B), with the following gradient: 10% B (2 min), 90% B (45 min), 90% B (50 min). Mass analysis was performed with a 150-1000 *m/z* scan at a resolution of 120.000. The area of the peaks of interest was measured and normalized on the signal of DMS (internal standard). Metabolites were harvested in 80% methanol with isotopically labeled amino acids and stored at -80 °C until analysis by LC-MS/MS.

### Western blotting

Larvae collected at 120 hpf (n=60) in triplicates were lysed in M-PER buffer (Cell Signaling Technology Inc., Danvers, MA) and Western blotting was performed as described elsewhere [20]. The following antibodies were used: anti-LC3 (NB100-2220, Novus Biologicals, Centennial, CO; 1:1000), anti-LAMP1 (ab24170, Abcam, Cambridge, UK, 1:500), anti-β-actin (GTX629630, GeneTex, Irvine, CA; 1:2500), and anti-vinculin (ab91459, Abcam, Cambridge, UK, 1:10000).

### RNA-Seq Analysis

The whole transcriptomic analysis was carried out on zebrafish 5dpf *cln5*^*−/−*^ and WT larvae. The WT larvae were also analyzed as a reference (Ctrl). Total RNA was extracted from larvae at 5 dpf (*n*= 36 larvae per group) using the Quick RNA miniprep kit (ZymoResearch, Irvine, CA), according to the manufacturer’s instructions, and checked for purity using NanoPhotometerTM Pearl, version 1.2 (IMPLEN, Westlake Village, CA); integrity (RNA integrity number > 7) was assessed using the RNA 6000 Pico Kit on a Bioanalyzer 2100 (Agilent Technologies, Santa Clara, CA). Indexed cDNA libraries were prepared from 350 ng of total RNA using the TruSeq Stranded kit (Illumina, San Diego, CA), quantified by real-time PCR, pooled at equimolar concentration, and sequenced with Illumina technology applying standard manufacturer protocols. The quality of reads was assessed using FastQC software Version 0.11.9 (http://www.bioinformatics.babraham.ac.uk/projects/fastqc/). Raw reads with more than 10% of undetermined bases or more than 50 bases with a quality score < 7 were discarded. Subsequently, reads were clipped from adapter sequences using Scythe software Version 0.994 (https://github.com/vsbuffalo/scythe), and low-quality ends (Q score < 20 on a 15-nt window) were trimmed with Sickle (https://github.com/vsbuffalo/sickle). Filtered reads were aligned to the current zebrafish reference genome assembly (GRCz11) using the STAR aligner (http://code.google.com/p/rna-star).

### Proteomic analysis

#### Sample preparation, proteolytic digestion, nano LC-MS/MS and data analysis

Zebrafish brains, eyes, and whole embryos were separately homogenized by adding a lysis buffer containing 8M urea (Fisher Chemical, Germany), 0.1M ammonium bicarbonate (ThermoFisher, Germany), 1% sodium deoxycholate (Thermo scientific, Italy), 0.1% Octyl α-D -glucopyranoside (Fluka, USA), and using Precellys®24 tissue homogenizer (Bertin Technologies, France). Proteins were reduced and alkylated using tris (2-carboxyethyl) phosphine (Aldrich, USA) and iodoacetamide (Sigma-Aldrich, USA) to a final concentration of 5 mM and 50 mM, respectively, and an incubation in the dark for 30 min. Ten micrograms of protein amounts were washed 6 times with 8M urea, 0.1M ammonium bicarbonate in Amicon Ultra-0.5 centrifugal filters (Millipore, Merck KGaA, Germany) using a modified FASP method [30,31]. The peptides were separated by Ultimate 3000 LC system (Dionex, Thermo Scientific) equipped with a reverse-phase trapping column RP-2TM C18 trap column (0.075 × 10mm, Phenomenex, USA), followed by analytical separation on a bioZen C18 nano column (0.075 × 250 mm, 2.6 μm particles; Phenomenex, USA). The injected sample analytes were trapped at a flow rate of 5 μl x min-1 in 100% of solution A (0.1% formic acid). After trapping, the peptides were separated with a linear gradient of 125 min comprising 110 min from 3% to 35% of solution B (0.1% formic acid/80% acetonitrile), 6 min 45% B, and 4 min to 95% of solution B. Each sample run was followed by at least one empty run to reduce the sample carryover from previous runs. LC-MS acquisition data was done with the mass spectrometer (Thermo Q Exactive HF) settings as follows: The resolution was set to 120,000 for MS scans, and 15000 for the MS/MS scans. Full MS was acquired from 350 to 1400 *m/z*, and the 15 most abundant precursor ions were selected for fragmentation with 45 s dynamic exclusion time. Ions with 2+, 3+, 4+, and 5+ charges were selected for MS/MS analysis. Maximum IT were set as 50 and 80 ms and AGC targets were set to 3e^6^ and 1e^5^ counts for MS and MS/MS respectively. Secondary ions were isolated with a window of 1 *m/z* unit. Dynamic exclusion was set with a duration of 45 s. The NCE collision energy stepped was set to 28 kJ mol–1. Following LC-MS/MS acquisition, raw files were qualitatively analyzed by Proteome Discoverer (PD), version 2.5 (Thermo Scientific, USA). The identification of proteins by PD was performed against the zebrafish protein databases (NCBI and SwissProt, downloaded in June 2023 with 45300 and 3277 entries, respectively) using the built-in SEQUEST HT engine. The following parameters were used: 10 ppm and 0.02 Da were tolerance values set for MS and MS/MS, respectively. Trypsin was used as the digesting enzyme, and two missed cleavages were allowed. The carbamidomethylation of cysteine residues was set as a fixed modification, while the oxidation of methionine, deamidation of asparagine and glutamine, and phosphorylation of serine and threonine were set as variable modifications. The false discovery rate was set to less than 0.01 and a peptide minimum length of six amino acids. Label-free quantification was done using unique peptides in Precursor Ion Quantifier.

#### Bioinformatic Analysis and Categorization of Data

Zebrafish brain and eyes were collected at different developmental stages, including 6 hpf and adult one. Samples were processed, as described elsewhere [32] and used for data dependent acquisition proteome analysis DDA. The NCBI and Swiss-Prot protein identifiers were further curated by adding the knowledge-based zebrafish IDs (ZFIDs, https://zfin.org/). Differentially expressed proteins (DEPs) were identified based on the number of unique peptides used for label-free quantitation (≥2), with an FDR < 0.01 and a fold change (FC) from averaged, normalized protein intensities diverging by >20%, both in up- and in down-regulation, and p ≤ 0.05 by ANOVA, in *cln5*^*−/*−^ versus the control group. Bioinformatic analysis was performed using the Ingenuity Pathway Analysis software platform (version: 111725566, release number: summer release, Q2 2024—June 2023; QIAGEN, Hilden, Germany) [33].

To assess gene expression, the normalized expression values for each transcript were calculated as Fragments Per Kilobase per Million mapped reads (FPKM). The expression profile of mutant *cln5*^*−/−*^ larvae was compared to the profile of WT and for each gene the ratio between average FPKM was calculated and reported as log2 fold change (log2FC). Transcripts showing a FC |≥2| and a False Discovery Rate (FDR, q-value) ≤ 0.01 were considered to estimate the predicted pathway activation or inhibition, and graphically represented.

To recognize the meaningful biological processes and molecular pathways, we performed a *Core Analysis* using the curated information from the QIAGEN Knowledge Base. The strength of the correlation between a subset of differentially expressed genes / proteins (DEGs/DEP) and a given biological function or disease was expressed by the p-value (calculated using a right-tailed Fisher’s Exact Test with Benjamini and Hochberg correction), whereas a “z-score” predicted the activation or inhibition of the given biological context. The macro-categories of Lipid Metabolism and Metabolic Disease were further scrutinized.

### Glucose uptake assay

An adapted Glucose Uptake-Glo™ Assay (Promega, Madison, WI) was performed on 96 hpf zebrafish to assess differences in the rate of glucose uptake between experimental groups, and following the already available protocol [34]. The protocol was adapted using a pool of 10 larvae for each experimental group. Moreover, we performed also an *in vivo* glucose uptake assay by using 2-deoxy-2-[(7-nitro-2,1,3-benzoxadiazol-4-yl)amino]-D-glucose (2-NBDG; Cayman Chemical Ann Arbor, MI). The larvae at 5dpf were incubated with 2-NBDG (600 μM) in E3 water for 3 h. After incubation, the larvae were rinsed with E3 water, using 0.04% tricaine anesthesia larvae were imaged using a using a Leica M205FA stereo-microscope (Leica Microsystems, Wetzlar, Germany), and fluorescence analysis was quantified using Image J v.1.46, calculating the fluorescent intensity in the region of interest (ROI).

### Statistics

All data in the manuscript represent three or more independent experiments. Statistical analysis was performed using GraphPad Prism 6. All quantitative variables were analyzed using either parametric or non-parametric methods, depending on the distribution shown by the Shapiro-Wilk test. For comparisons between two different groups the analysis was performed using the T-test for normally distributed data; instead, the Mann-Whitney test was applied in the case of non-normally distributed data. For multiple comparisons, Dunn’s test was performed after the Kruskal-Wallis test, since the data of the various groups examined did not show Gaussian distribution. Statistical significance is reported as follows: ^*^ *p* ≤ 0.05, ^**^ *p* ≤ 0.01, ^***^ *p* ≤ 0.001, or ^****^ *p* ≤ 0.0001. The specific test applied for each analysis is reported in the figure legend.

## Results

### Generation and characterization of novel *cln5*^*−/−*^ zebrafish line

We evaluated the spatiotemporal expression of *cln5* during zebrafish development. *In situ* hybridization of *cln5* mRNA revealed that it is expressed over the entire larval body at 5 dpf with highest expression at brain level (**Figure 1A**). We found it to be expressed from the first stage of development (it is highly expressed at 0 hpf); after midblastula transition at 10 hpf its expression decreases and thereafter it is constantly expressed until 5 dpf (**Figure 1B**). We successfully edited the novel *cln5*^*−/−*^ zebrafish model using the CRISPR/Cas9 system, which resulted in a 2-bp deletion and a missense mutation in exon 2. This led to a frameshift mutation and an early stop codon at residue 69 (**Figure 1C**). The efficacy of the mutation was further confirmed by qRT-PCR analysis in homozygous F2 mutant *cln5*^*−/−*^ larvae, which showed reduced expression of *cln5* mRNA versus control siblings (**Figure 1D**), likely as a consequence of nonsense-mediated mRNA decay (NMD). We further confirmed gene depletion by performing mass spectrometry assessment of BMP production, detecting a high decrease of (18:1/18:1) BMP (S,S) in *cln5*^*−/−*^ mutant larvae as compared to WT controls (**Figure 1E**). The mutation did not affect viability, and the Mendelian ratios were maintained. We also observed no qualitative difference in the phenotype between zygotic *cln5*^*−/−*^ embryos derived from heterozygous females and maternal-zygotic (MZ) *cln5*^*−/−*^ embryos obtained from homozygous mutant females. This finding suggests that maternal *cln5* mRNA does not play a crucial role during early embryonic development in zebrafish. Mutant embryos and larvae demonstrated an altered behavior associated with reduced locomotor performances in distance and velocity travelled in larvae (**Figure 1F**). Since Purkinje cells (PCs) are the most affected neuronal cells in the brain of NCL patients and animal models [35,36], we used the transgenic strain *Tg(aldoca:GAL4FF)* to analyze the PCs, and observed decreased PCs layer compared to WT siblings (**Figure 1G**).

**Figure 1.**
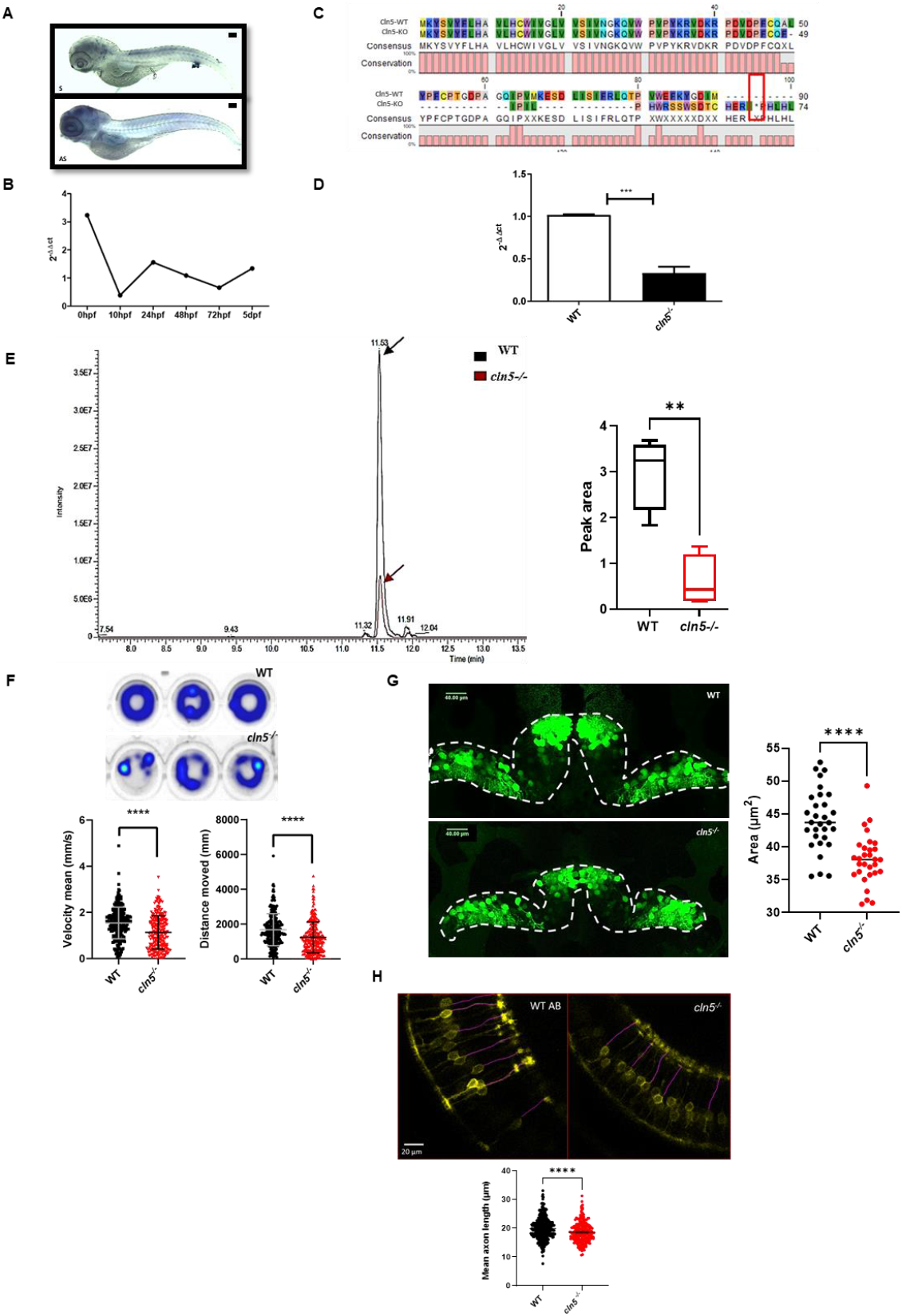
Characterization of the novel *cln5*^*−/−*^ zebrafish mutant line. (**A**) Whole mount in situ hybridization of the *cln5* transcript in 5 dpf larvae, showing the sense probe (S) and the antisense probe (AS). (scale bar=100μm). (**B**) Temporal expression of *cln5* mRNA at different developmental stages using q-RT-PCR. (**C**) Protein alignment showing the early stop codon generated by CRISPR/cas9 system in mutant larvae (*cln5*^*−/−*^) at position 69 (p.Ala49Phefs*69) compared to WT siblings. (**D**) The graph represents the decreased expression of *cln5* mRNA in mutant *cln5*^*−/−*^ larvae at 5dpf compared to WT. The analysis was performed in triplicate using n=30 for each experimental group. Statistical analysis has been performed using t-test (^***^ P≤ 0.001). (**E**) LC/MS dosage of BMP in adult brain of *cln5*^*−/−*^ mutant (red line and arrow; n=4) and WT zebrafish (black line and arrow; n=4), showing a decreased BMP production in *cln5*^*−/−*^, the graph shows the statistical analysis performed using t-test (^**^ P ≤ 0.01). Areas are normalized on the signal of N,N-dimethylsphingosine (internal standard) (**F**) Locomotion analysis of 5 dpf WT and *cln5*^*−/−*^ larvae. (**G**) Confocal images of the Purkinje cell layer, the graphs show the analysis of the area of the PCs layer in 5 dpf WT (*n*=30) and *cln5*^*−/−*^ (*n*=30) larvae. All data were obtained from three independent experiments and were analyzed through statistical analysis (^****^ P≤ 0.0001) performed using the Mann-Whitney test. The values are expressed as mean ± standard error of the mean (SEM). (**H**) Analysis of axons length of retinal ON bipolar cells using confocal imaging, the graph shows a decreased length in *cln5*^*−/−*^ (*n*=39) compared to WT (*n*=45), data were analyzed through statistical analysis (**** P ≤ 0.0001) performed using the Mann-Whitney test. All data were obtained from three independent experiments. The values are expressed as mean ± standard error of the mean (SEM).

To further confirm the retinal dysfunction we used the transgenic line *Tg(nyx:GAL4-VP16)* tagging retinal ON-bipolar cells, and showed that *cln5* depletion causes a decrease in axonal length of retinal cells (**Figure 1H**).

### Transcriptomic analysis revealed impaired lipid and glucose metabolisms

RNA-seq analyses showed 76 upregulated and 101 downregulated genes in *cln5*^*−/−*^ larvae compared to WT (**Figure 2A**). Canonical pathway analysis using Ingenuity Pathway Analysis (IPA) revealed that most of the DEGs in mutants compared to WT were associated with apoptosis, visual function and cell cycle, while the most downregulated pathways were linked to glucose metabolism, such as the regulation of insulin growth factor (IGF) transport and uptake, glycolysis and gluconeogenesis (**Figure 2B**). IPA surveys revealed that *cln5*^*−/−*^ larvae exhibited a decrease in glucose uptake and impaired glucose metabolism (**Figure 2C**).

**Figure 2.**
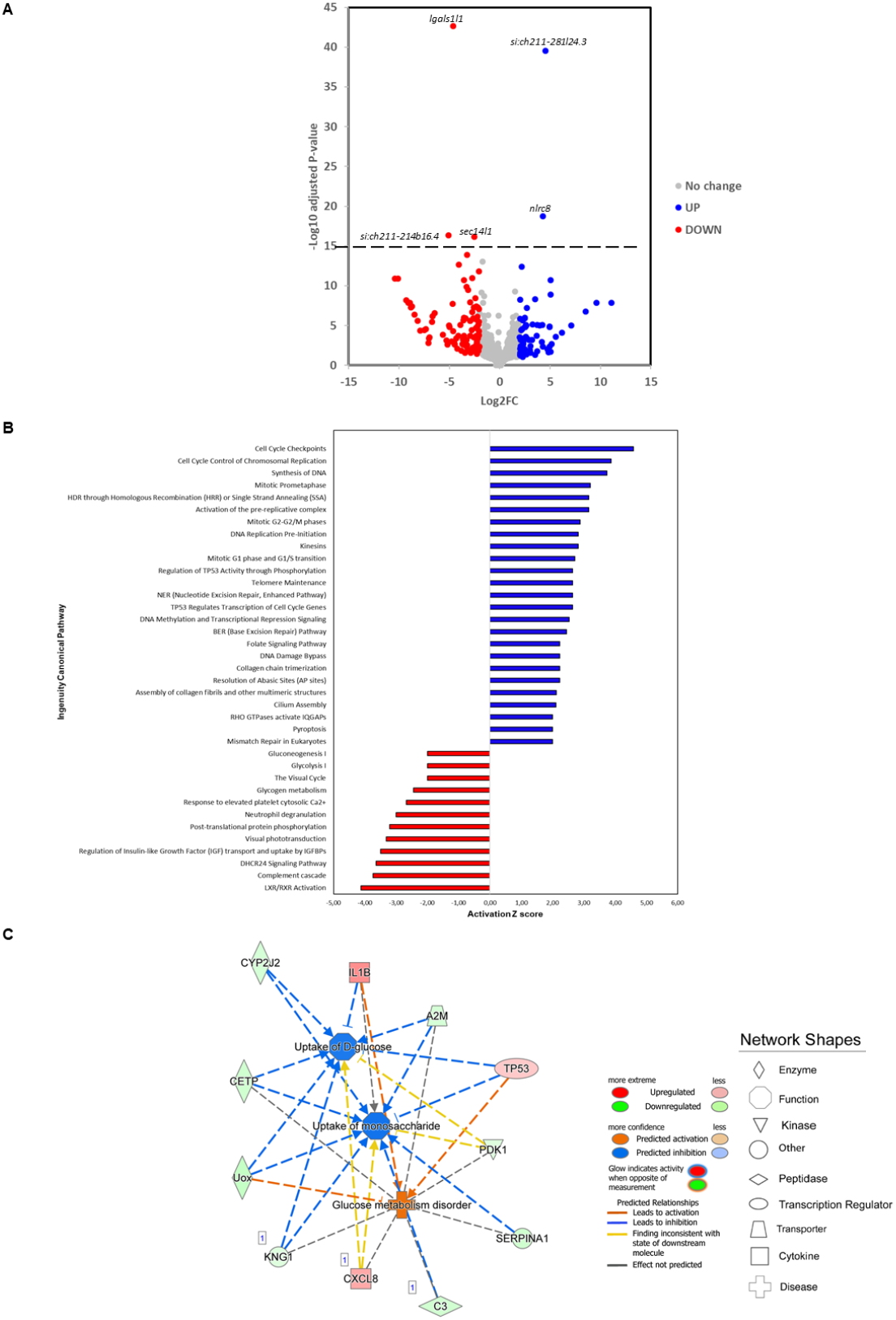
Transcriptomic analyses of whole larvae. **(A)** Volcano plot. The log2 FC indicates the mean expression level for each gene. Each dot represents one gene. The names near the dots indicate the genes with values with a log10 of adjusted p-value major than 15, the dotted line represents a threshold settled at 15. **(B)** RNA-seq analysis performed on 5 dpf larvae, the graph showed the canonical pathways altered in *cln5*^*−/−*^ mutant compared to WT. The analysis has been performed using maximal stringency with p value <0.01 and z score >2 and <-2. **(C)** Two distinct networks were reconstructed to be involved in Glucose Metabolism (Focused Molecules *n*= 11).

### Proteomic analysis at different developmental stages of *cln5*^*−/−*^ mutant zebrafish

Proteomic analyses at different developmental stages (namely, 6 hpf and adult brain and eyes) showed several DEPs in our *cln5*^*−/−*^ samples compared to WT. IPA analyses on adult brain and eye samples, confirmed RNA-seq findings with altered lipid metabolism (**Figure 3A**) and altered glucose metabolism (**Figure 3B**), in both the brain and eyes being the most statistically significant. To confirm the results on altered glucose metabolism we performed a glucose uptake assay, which demonstrated a profound decrease in glucose uptake in *cln5*^*−/−*^ larvae compared to the WT ones (**Figure 3C-D**), thereby functionally validating the transcriptomic and proteomic data.

**Figure 3.**
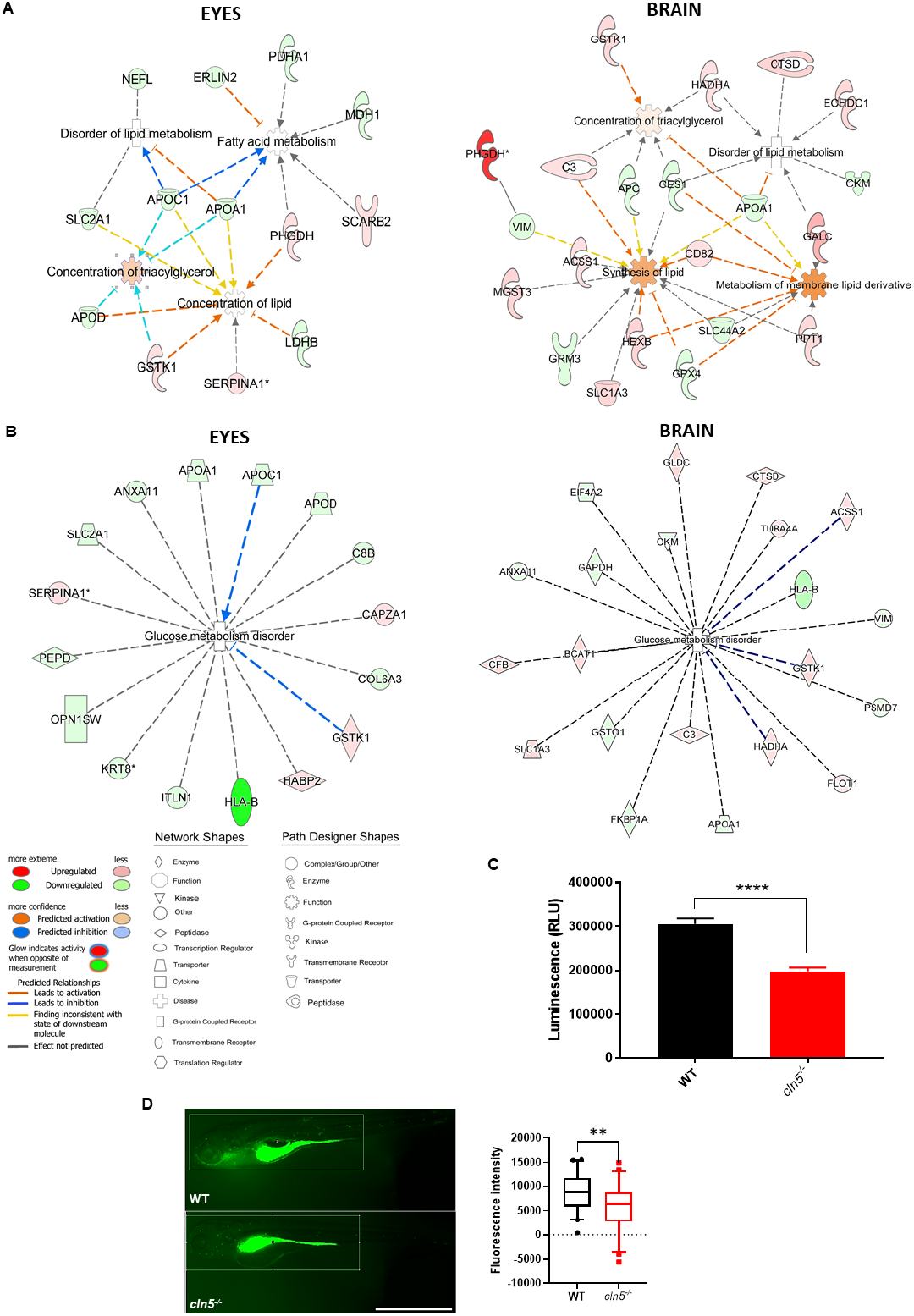
Proteomic analyses of isolated brains and eyes of adult fish using IPA software. (**A**) The networks in both brain and eyes show alterations in lipid metabolism. (**B**) The networks portraying the DEPs involved in altered glucose metabolism. (**C**) The graph represents the glucose uptake assay in *cln5*^*−/−*^ and WT 96 hpf larvae. The values are expressed as mean ± standard error of the mean (SEM). Statistical analysis was performed using the Mann-Whitney test (^****^ P ≤ 0.0001). (**D**) *In vivo* glucose uptake analysis using 2-NBDG, figure show images of zebrafish larvae at 5 dpf (scale bar 1000 μm), the rectangle indicate the ROI used for the analysis of fluorescence intensity, the graph showed the quantification of fluorescence intensity in n=45 *cln5*^*−/−*^ and WT respectively. The values are expressed as mean ± standard error. Statistical analysis was performed using the Mann-Whitney test (^**^ P ≤ 0.01). All data were obtained from three independent experiments.

## Discussion

The limited understanding of CLN5-related cellular mechanisms and lack of defined pharmacological targets underscore the need to study the developmental impact of CLN5 loss, with a focus on uncovering pathways and biomarkers relevant to juvenile dementia [2]. Our novel zebrafish *cln5*^*−/−*^ model recapitulates the main features of the late-infantile form of Batten disease, including decreased BMP synthesis [11], increased cell death, accumulation of subunit C of ATP synthase, “ataxia-like” motor behavior and vision loss. Moreover, in-depth characterization also revealed retinal degeneration and decreased axonal length [37], in ON-bipolar cells, resembling the dysfunction already observed in the CLN5 KO ovine model [38]. These parallels make the *cln5*^*−/−*^ zebrafish a powerful tool for studying the neurodegenerative progression in CLN5 disease, especially in early-life stages, to better understand the cellular changes contributing to juvenile dementia. Interestingly, a decreased PC layer has also been observed upon *cln5* depletion, a finding common to other NCLs forms [36], which is further supported by an increased apoptosis in the brain at early stages (24 hpf). Taken together, our gene editing and follow-up characterization in zebrafish offers a new tool to investigate CLN5-related pathology during early development. As observed in similar NCL models [20,39], zebrafish strains are particularly amenable for drug screening or repurposing. Three additional points of interest emerged from the current study.

We performed multi-omics studies in larvae and adult brains and eyes disclosing, at different developmental stages (embryos and adults) the same differentially abundant proteins (i.e. Ctsd, Hexb), already pinpointed by lysosomal-targeted organelle proteomics in SHSY-5Y KO *CLN5* [12]). These data point to the already observed lysosomal-autophagy balance and lipid metabolism [40,12], suggesting the zebrafish KO strain to be sufficiently robust to model the human disease status. Original data pointed to alteration in glucose metabolism with decreased uptake of glucose, which has been assorted to *cln5* dysfunction for the first time, though a hint to this information had been partially presented by others [41,42]. Furthermore, we observed a predicted downregulation of the pathway related to insulin growth factor 1 (IGF1) transport and uptake, which can be considered as a potential drug target for CLN5 and other NCLs forms, as seen in *mnd/mnd* mouse model [43]. This metabolic disruption resembles energy imbalances reported in other neurodegenerative diseases and could underlie dementia-related symptoms such as memory loss and executive dysfunction. Further research is required to clarify these potential links and to explore therapeutic strategies targeting glucose metabolism in NCLs.

In summary, the development of this innovative *in vivo* model of CLN5 offers a groundbreaking platform that faithfully mimics the behavioral, physiological, and molecular characteristics of CLN5 disease. This model not only opens new avenues for exploring the novel functions of the CLN5 protein in axonal growth and also sheds light on an altered glucose metabolism linked to CLN5 loss-of-function. Such insights could lead to exciting new strategies to halt or delay the progression of CLN5-related human disease.

## Ethics statement

The generation of the mutant, using CRISPR/Cas9 technology, was performed with the ethical approval of the Italian Ministry of Health (approval n° 584/2019-PR), in accordance with the European Union’s Directive 2010/63/EU on the protection of animals used for scientific purposes, under the supervision of the University of Pisa Animal Care and Use Committee, and in compliance with the 3R principles.

## Acknowledgements

We are grateful to Dr. Baldassare Fronte of the University of Pisa for his technical support in the management of the zebrafish facility. The authors thank the Italian patients’ association (A-NCL) for their constant encouragement and support. Proteomics measurements were performed at the Meilahti Clinical Proteomics Core Facility (supported by HiLIFE and Biocenter Finland).

## Contributions

Conceptualization: M.M.; Methodology: M.M. S.B., R.V.; S.D.; L.S; R.S.; S.M.; R.L; A.O. Formal analysis and investigation: M.M.; S.B.; R.V.; S.D.; L.S; M.L.; S.M.; R.L; J.Z; G.S. Writing - original draft preparation: M.M.; Writing - review and editing: F.M.S.; S.B.; S.D.; M.L.; S.M.; R.L; J.Z.; S.N.; R.S.; G.S.; Funding acquisition: M.M. and F.M.S. Resources: M.M. and F.M.S.; Supervision: M.M. and F.M.S.

